# Hippocampal GFAP-positive astrocyte responses to amyloid and tau pathologies

**DOI:** 10.1101/2022.02.25.481812

**Authors:** Marco Antônio De Bastiani, Bruna Bellaver, Wagner S. Brum, Debora G. Souza, Pamela C. L. Ferreira, Andreia S. Rocha, Guilherme Povala, João Pedro Ferrari-Souza, Andrea L. Benedet, Nicholas J. Ashton, Thomas K. Karikari, Henrik Zetterberg, Kaj Blennow, Pedro Rosa-Neto, Tharick A. Pascoal, Eduardo R. Zimmer, the Alzheimer’s Disease Neuroimaging Initiative

## Abstract

**Introduction:** In Alzheimer’s disease clinical research, glial fibrillary acidic protein (GFAP) released into the cerebrospinal fluid and blood is widely measured and perceived as a biomarker of reactive astrogliosis. However, it was demonstrated that GFAP levels differ in individuals presenting with amyloid-β (Aβ) or tau pathology. The molecular underpinnings behind this specificity are unexplored. Here we investigated biomarker and transcriptomic associations of GFAP-positive astrocytes with Aβ and tau pathologies in humans and mouse models.

**Methods:** We studied 90 individuals with plasma GFAP, Aβ- and Tau-PET to investigate the association between biomarkers. Then, transcriptomic analysis in hippocampal GFAP-positive astrocytes isolated from mouse models presenting Aβ (PS2APP) or tau (P301S) pathologies was applied to explore differentially expressed genes (DEGs), Gene Ontology processes, and protein-protein interaction networks associated with each phenotype.

**Results:** In humans, we found that plasma GFAP associates with Aβ but not tau pathology. Unveiling the unique nature of GFAP-positive astrocytic responses to Aβ or tau pathology, mouse transcriptomics showed scarce overlap of DEGs between the Aβ and tau mouse models, While Aβ GFAP-positive astrocytes were overrepresented with genes associated with proteostasis and exocytosis-related processes, tau hippocampal GFAP-positive astrocytes presented greater abnormalities in functions related to DNA/RNA processing and cytoskeleton dynamics.

**Conclusion:** Our results offer insights into Aβ- and tau-driven specific signatures in GFAP-positive astrocytes. Characterizing how different underlying pathologies distinctly influence astrocyte responses is critical for the biological interpretation of astrocyte-related biomarker and suggests the need to develop context-specific astrocyte targets to study AD.

**Funding:** This study was supported by Instituto Serrapilheira, Alzheimer’s Association, CAPES, CNPq and FAPERGS.

## 1. Introduction

Astrocytes are the main homeostatic cells of the brain, which are involved in most aspects of brain physiology. For a long time, the response of astrocytes to brain injury – a process collectively called astrocyte reactivity – was considered an almost universal phenomenon. Nowadays, there is a consensus that glial responses are rather complex, heterogeneous, and context-specific [1–3]. In this sense, astrocytes are likely decisive contributors to Alzheimer’s disease (AD) onset and progression; however, the molecular underpinnings are still poorly understood. Furthermore, transcriptomics analysis in *postmortem* AD brains demonstrated highly variable intra-cohort astrocyte molecular signatures in individuals with the same clinical diagnosis, indicating that specific aspects of AD pathophysiology might be triggering different astrocyte phenotypes [4]. In this context, Jiwaji and colleagues started uncovering the heterogeneity of astrocyte reactivity facing Aβ and tau pathologies, the neuropathological hallmarks of AD [5].

Astrocyte biomarkers are reported to be consistently increased in cerebrospinal fluid (CSF) and blood of AD patients compared to cognitively unimpaired (CU) individuals [6, 7]. Although it is known that astrocyte reactivity cannot be measured by one single marker, clinical research commonly relies on the measurement of glial fibrillary acidic protein (GFAP) concentration in CSF or blood as a proxy of astrocyte reactivity. It was recently demonstrated that plasma GFAP levels are more associated with the Aβ burden than tau pathology in AD [8, 9] – an association that was found to be more robust in blood than in CSF. On the other hand, increased GFAP levels were already identified in primarily tau pathologies as in frontotemporal dementia [10–12]. In this sense, uncovering the phenotypes acquired by GFAP-positive astrocytes under Aβ or tau influence might directly impact clinical research by improving the biological interpretation of clinical biomarkers used in AD.

In this study, we investigated GFAP associations with Aβ and tau pathologies in individuals across aging and AD spectrum. Furthermore, we investigated the molecular signatures of GFAP-positive astrocytes driven by Aβ or tau, comparing the hippocampal transcriptomic profile of astrocytes from animal models of amyloidosis (PS2APP mice) and tau pathology (P301S mice). Differentially expressed genes (DEGs), protein-protein networks, and functional enrichment analysis of Gene Ontology (GO) terms were performed to identify biological differences and similarities between the phenotypes acquired by these astrocytes. The molecular characterization of astrocyte responses to Aβ and tau pathology may pave the way for a better understanding of the GFAP biological interpretation in AD.

## 2. Methods

### 2.1. Human Cohort

Data used in the preparation of this article were obtained from the Alzheimer’s Disease Neuroimaging Initiative (ADNI) database, phases GO and 2. ADNI was launched in 2003 as a public-private partnership, led by Principal Investigator Michael W. Weiner, MD. The primary goal of ADNI has been to test whether serial magnetic resonance imaging (MRI), PET, other biological markers, and clinical and neuropsychological assessments can be combined to measure the progression of mild cognitive impairment (MCI) and early AD. ADNI study was conducted according to Good Clinical Practice guidelines, US 21CFR Part 50 – Protection of Human Subjects, and Part 56 – Institutional Review Boards, and pursuant to state and federal HIPAA regulations and was approved by the Institutional Review Board of each participating site (adni.loni.usc.edu). Written informed consent for the study was obtained from all participants and/or authorized representatives

Based on biomarker availability, 90 participants were included [CU: n = 39; cognitively impaired (CI; mild cognitive impairment (MCI) or AD dementia): n = 51] from the ADNI database. All participants had available measurements of plasma GFAP and positron emission tomography (PET) data for [^18^F]-florbetapir (Aβ-PET) and [^18^F]-flortaucipir (Tau-PET), within a maximum 2.5-year time interval between scan and plasma collection. Plasma GFAP was measured with Simoa Neurology 4-Plex (#103670, Quanterix). For Aβ-PET data analysis, as recommended for cross-sectional analyses [13], the global cortical composite standardized uptake value ratio (SUVr) normalized by the whole cerebellum was used. When applicable, global Aβ-PET cortical composite was used to dichotomize participants into Aβ-negative (A-) and Aβ-positive (A+) based on the widely validated ADNI cutoff of >1.11 [13]. For tau-PET, we assessed the temporal meta-ROI values and the cutoff for tau-PET positivity of 1.23 was used [14]. Baseline characteristics were summarized with descriptive statistics. Pearson correlation was applied to assess associations between biomarkers, and statistical significance was set to α = 0.05 (FDR-adjusted).

### 2.2. RNA-seq Data Acquisition and Differential Expression Analysis

Expression profiling by high throughput sequencing studies of mice hippocampus GFAP-positive astrocytes were obtained from the NCBI Gene Expression Omnibus (GEO) (https://www.ncbi.nlm.nih.gov/geo). The first study (GSE129770) contains data from 5 PS2APP and 5 wild-type (WT) mice (11.5-months old); the second study (GSE129797) contains data from 5 Tau P301S and 5 WT mice (6-months old)[15]. First, raw RNA-seq data from both studies was downloaded using the SRA Toolkit (https://github.com/ncbi/sra-tools). Afterwards, transcript alignment was performed using Salmon (v1.3.0) [16], mapped to a reference genome with the index derived from *Mus musculus* GRCm38 Ensembl build. Aligned reads were summarized using tximport (v1.12.3) [17] and genes with mean count < 10 were filtered out. Finally, processed expression data from both hippocampus astrocyte studies was analyzed with the DESeq2 (v1.28.1) [18] method for differential expression evaluation of either PS2APP or Tau P301S mutations *versus* respective WT mice. Genes with FDR adjusted p-value < 0.1 were considered as DEGs for further analyses. Venn diagrams were constructed using the VennDiagram package (v1.6.20).

### 2.3. Functional Enrichment Analysis and Network

Gene Ontology (GO) enrichment analysis of DEGs previously obtained for both studies was implemented using the clusterProfiler (v3.16.1) package [19]. The significantly enriched terms from biological process (BP), cellular component (CC) and molecular function (MF) sub-ontologies were used to build networks employing the RedeR (v1.36.0) and GOplot (v1.0.2) packages [20, 21]. In this network, GO terms are mapped to nodes and edges represent the proportion of gene overlap (Jaccard Coefficient > 0.15) between all pairs of terms.

### 2.4. Protein-Protein Interaction Networks and Circos Plots

We used the *Search Tool for the Retrieval of Interacting Genes/Proteins* (STRING) Consortium to build protein-protein interaction (PPI) networks with the DEGs for both datasets. STRING is a biological database and web resource of known and predicted protein–protein interactions which contains information from numerous sources, including experimental data, computational prediction methods and public text collections. The construction of the networks was implemented in R using the STRINGdb (v2.0.2), RedeR (v1.36.0) and igraph (v1.2.6) packages [20, 22]. Final network was filtered to retain only gene connections with STRING combined confidence interaction scores > 0.7. For each PPI network, genes observed in GO terms significantly enriched were manually curated and categorized. These functional categorizations were used to create circos plots showing category linkages and link proportions in PS2APP and Tau P301S networks. The plots were built using the circlize package (v0.4.13) [23].

## 3. Results

### 3.1. Plasma GFAP differently associates with biomarkers of AD pathology

**Table 1** shows demographic information of the 90 individuals (39 CU and 51 CI) assessed. Correlations between GFAP and other known biomarkers of AD pathology and neurodegeneration showed that GFAP negatively correlated with CSF Aβ42/40, but no significant associations were observed with p-Tau181 (**Supplemental Fig. 1**). We then further investigated associations between Aβ- and Tau-PET with plasma GFAP levels in individuals across aging and the AD continuum. Linear regressions using composite brain regions revealed that both Aβ- and Tau-PET significantly associated with plasma GFAP (**Fig. 1A, C**). However, when individuals were stratified by their Aβ status (A- and A+), we observed that the association between Aβ- and Tau-PET with plasma GFAP remains significant only in A+ individuals (**Fig. 1B, D**).

**Table 1.** Demographic informationFigure Legends

|  | CU<br>(N=39) | CI<br>(N=51) |
| --- | --- | --- |
| *Demographics* |  |  |
| Age, years, mean (SD) | 71.7 (6.07) | 71.5 (5.92) |
| Sex, female, n (%) | 21 (53.8%) | 19 (37.3%) |
| Education, years, mean (SD) | 16.7 (2.34) | 16.2 (3.04) |
| Baseline MMSE score, mean (SD) | 29.4 (0.72) | 26.3 (4.33) |
| APOE ε4 carriers, n (%) | 15 (38.5%) | 22 (43.1%) |
| *Biomarkers* |  |  |
| [$^{18}$F]-florbetapir global SUVR, mean (SD) | 1.11 (0.158) | 1.19 (0.200) |
| Aβ-PET-positive, n (%) | 14 (35.9%) | 30 (58.8%) |
| [$^{18}$F]-flortaucipir temporal meta-ROI SUVR, mean (SD) | 1.19 (0.092) | 1.29 (0.233) |
| Tau-PET-positive, n (%) | 10 (25.6%) | 26 (51.0%) |
| Plasma GFAP, pg/mL, mean (SD) | 145 (59.2) | 154 (79.5) |

**Figure 1.**
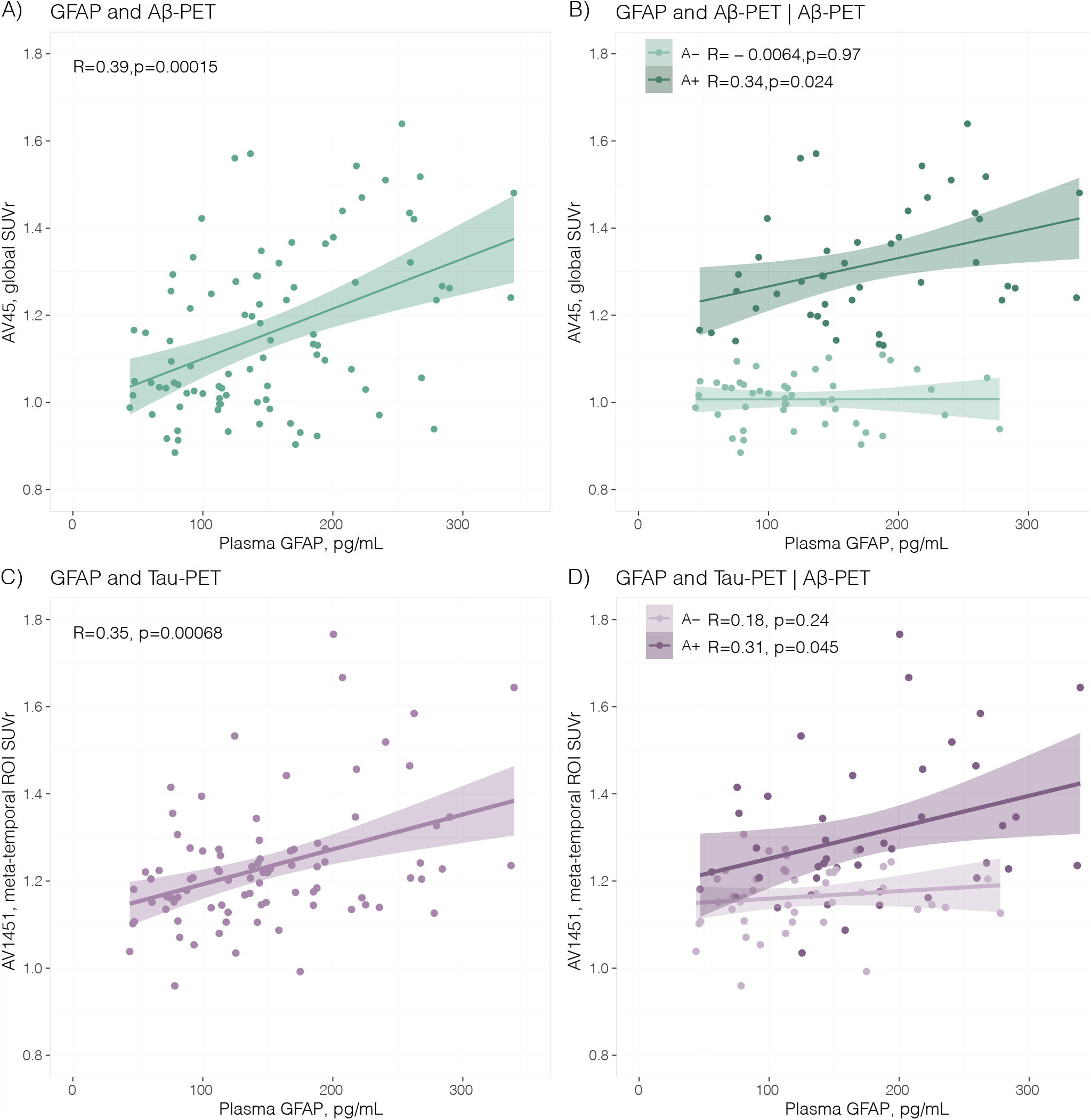
Associations of plasma glial fibrillary acidic protein with Aβ and Tau-PET. Association between plasma GFAP and **(A)** Aβ-PET ([^18^F]AV45) and **(C)** Tau-PET ([^18^F]AV1451). Association of plasma GFAP and with Aβ- and Tau-PET stratified by **(B)** Aβ and **(D)** tau positivity (n = 90).

Furthermore, we evaluated whether GFAP levels are associated with changes in Tau-PET, to further assess how this effect would be affected by including Aβ-PET as a covariate. We built models including either (i) Braak III-IV or (ii) Braak V-VI SUVr as the outcomes, with two predictor schemes, plasma GFAP alone or plasma GFAP and Aβ-PET global SUVr, resulting in four models. For Braak III-IV SUVr as the outcome, increases in plasma GFAP were associated with increases in Tau-PET (β-estimate: 0.054; t = 3.68; p < 0.001). When including plasma GFAP and Aβ-PET as predictors, plasma GFAP was no longer associated with increases in Tau-PET (β-estimate: 0.019; t = 1.35; p = 0.18). For Braak V-VI as the outcome, the same was observed. In the univariate model, increases in plasma GFAP were associated with increases in Tau-PET (β-estimate: 0.031; t = 2.43; p = 0.02). When Aβ-PET SUVr was also added as predictor, plasma GFAP was no longer associated with increases in Tau-PET (β-estimate: 0.0059; t = 0.46; p = 0.65). Our results corroborate previous studies in other cohorts [8, 9], evidencing the specificity of plasma GFAP association with Aβ, but not tau pathology in AD.

### 3.2. GFAP-positive astrocytes affected either by Aβ or tau pathology present a scarce overlap of DEGs

To better understand the biological underpinnings of GFAP association with Aβ pathology, we compared GFAP-positive astrocytes isolated from Aβ and tau mouse models.

The amyloidosis model (PS2APP) presented 95 downregulated genes (adjusted p-value < 0.1) compared to WT mice (**Fig. 2A**). Log fold change values ranged from −2.9365 to - 0.4204. The most significantly downregulated gene was Aquaporin 11 (Aqp11), which encodes a water channel protein that also facilitates the transport of glycerol and hydrogen peroxide across membranes. In addition, 200 upregulated genes were also found compared to WT mice (adjusted p-value < 0.1) comprising log fold change values ranging from 0.4308 to 4.9284. The most significant upregulated gene was PTPN6, which encodes the Protein Tyrosine Phosphatase Non-receptor Type 6, also known as Src homology region 2 domaincontaining phosphatase-1 (SHP-1), a tyrosine phosphatase enzyme. The Tau model (P301S) presented 178 statistically significant downregulated genes compared to WT mice (**Fig. 2B**). The expression of Insulin-1 (Ins1) and Insulin-2 (Ins2) were mostly impacted. These genes encode the hormone insulin, and may play a role in growth, differentiation and glucose metabolism in the brain. The other downregulated genes ranged between −4.098 to −0.507 log fold change and are related to different pathways. In parallel, 427 statistically significant upregulated genes were found in Tau P301S compared to WT mice. The log fold change values ranged from 0.5216 to 4.212. The most significant upregulated gene identified was Retrotransposon Gag Like 8B (Rtl8b), also known as Mart, considered a neofunctionalized retrotransposon gene. Interestingly, only 30 DEGs overlapped among astrocytes from these two mouse models, revealing Aβ and tau drive specific astrocyte molecular remodeling (**Fig. 2C**). Results of differential expression analysis for both animal models are presented in **Supplemental Table 1**.

**Figure 2.**
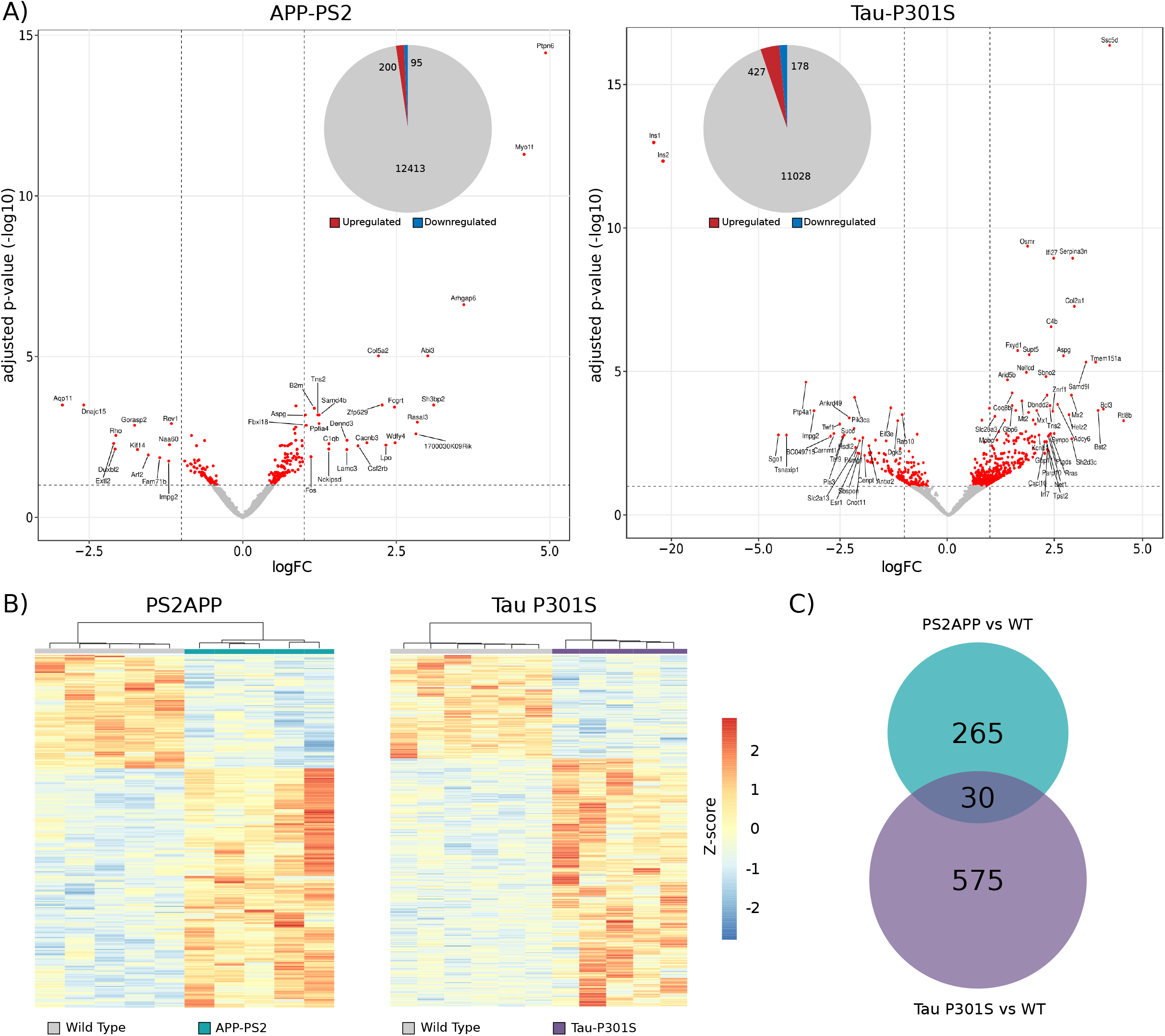
Gene expression analysis in hippocampal astrocytes from AD mouse models. **(A)** Volcano plots depicting differentially expressed genes (DEGs) in PS2APP (left) or Tau P301S (right) versus WT astrocytes. Insert shows a pie chart of upregulated and downregulated DEG counts. **(B)** Heatmaps with hierarchical clustering depicting DEGs differentiating PS2APP (left) or Tau P301S (right) versus WT astrocytes. **(C)** Venn diagram showing DEGs in each model and the matched genes altered in both.

### 3.3. Aβ and tau pathology promote the enrichment of distinct functional categories in GFAP-positive astrocytes

The PPI analysis showed a network containing 70 altered nodes in the PS2APP hippocampal astrocytes dataset. The average node degree was 3.285 and the number of edges was 115 (expected: 74). The average local clustering coefficient was 0.521, while the global clustering coefficient was 0.21. Based on the connectivity degree, Uba52, Ubc, Ccnh, Arrb2, and Gng12 were the top 5 hub proteins. The results were plotted as a PPI network (**Fig. 3A**). Circos plot (**Fig. 3B**) indicates the altered protein interconnectivity in the network based on their functional categories, with “Proteostasis” being the leading functional alteration in PS2APP astrocytes.

**Figure 3.**
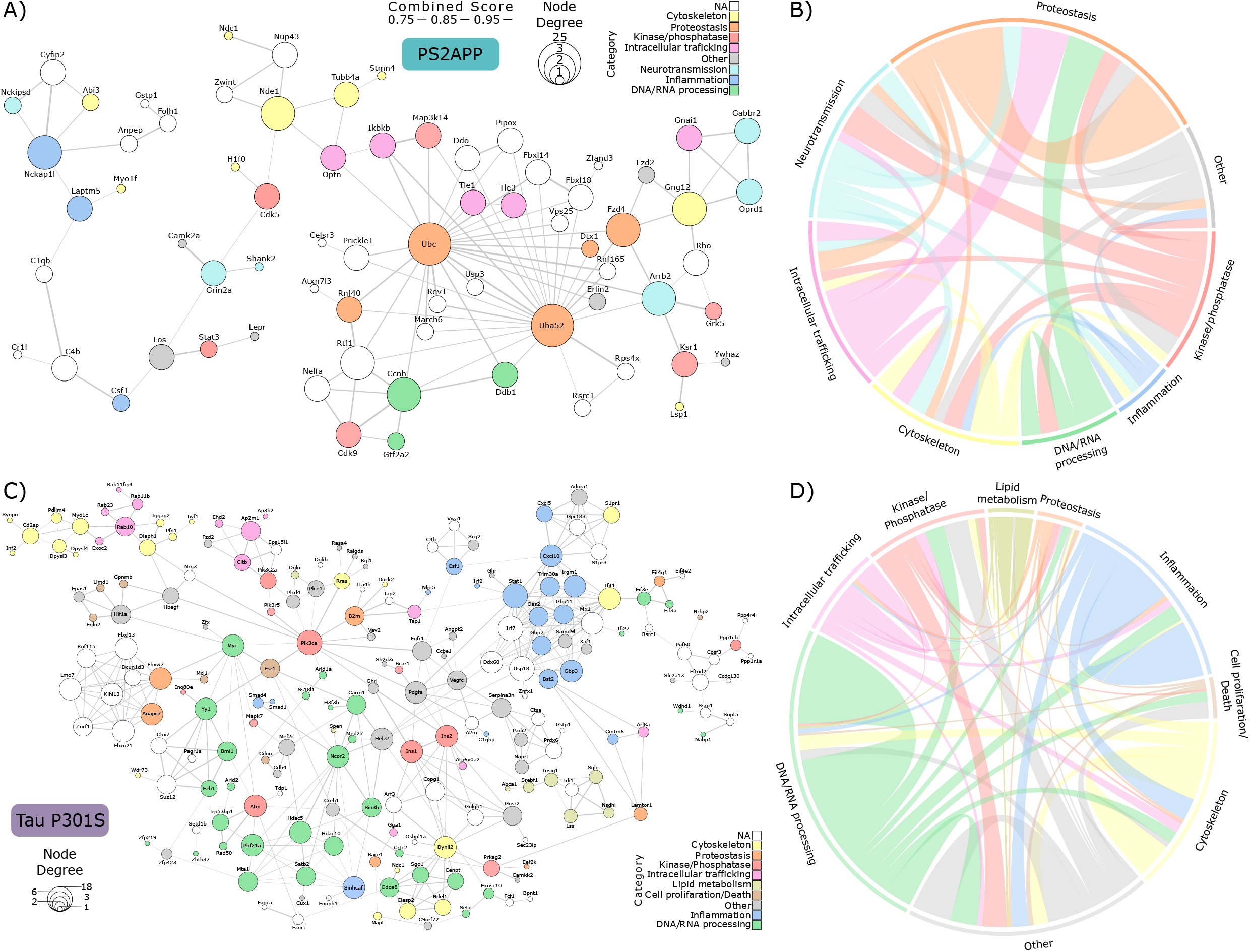
Protein-protein interaction (PPI) networks in PS2APP and Tau P301S astrocytes. PPI network of **(A)** PS2APP and **(C)** Tau P301S mapping functional categories of genes observed in enriched Gene Ontology biological processes (GOBP). “NA” denotes genes not observed in any enriched GOBPs. Combined score mapped to edges represents the combined confidence interaction scores provided by the STRING database for each pair of genes (values >0.7 are considered high confidence scores). Circos plots showing the connections between functional categories in **(B)** PS2APP and **(D)** Tau P301S PPI.

The PPI data analysis showed a different profile of the gene network in the Tau P301S model compared to the PS2APP. PPI analysis of the Tau P301S hippocampal astrocytes showed 204 altered nodes. The average node degree was 4.137 and the number of edges was 422 (expected: 373). The average local clustering coefficient was 0.551, and the global clustering coefficient was 0.5. Based on the connectivity degree, Stat1, Usp18, Irf7, Pik3ca, and Cxcl10 were among the top 5 hub proteins. The results were plotted as a PPI network and protein nodes were colored according to their functional category (**Fig. 3C**). Circos plot (**Fig. 3D**) indicates the interconnectivity of the altered proteins based on their functional categories, being “DNA/RNA processing”, “Inflammation” and “Cytoskeleton” the leading functional alterations. The modulation of gene expression for both networks can be visualized in **Supplemental Fig. 2**.

### 3.4. Aβ and tau pathologies drive GFAP-positive astrocytes changes in Gene Ontology biological domains

The functional enrichment analysis revealed 36 GO terms enriched with DEGs altered in the PS2APP mice compared to its WT controls (**Supplemental Table 2**). We found 14 BP, 8 CC and 14 MF in the analysis. Fig. 4A shows the network of GO terms enriched, in which we mapped the degree of connectivity for each term in the node sizes and the pairwise gene overlap (Jaccard coefficient) between terms in the edges. We noted that “exocytosis”, “exocytic process”, “protein localization to cell periphery”, “protein localization to plasma membrane” and “voltage–gated cation channel activity” are the most connected terms in the network (Fig. 4B). On the other hand, the most significantly enriched terms are shown in **Fig. 4C**, which include processes such as “protein localization to cell periphery and plasma membrane”, “interleukin-6 production and regulation”, “interleukin–6 production and regulation”, “exocytosis” and “neuron death”.

**Figure 4.**
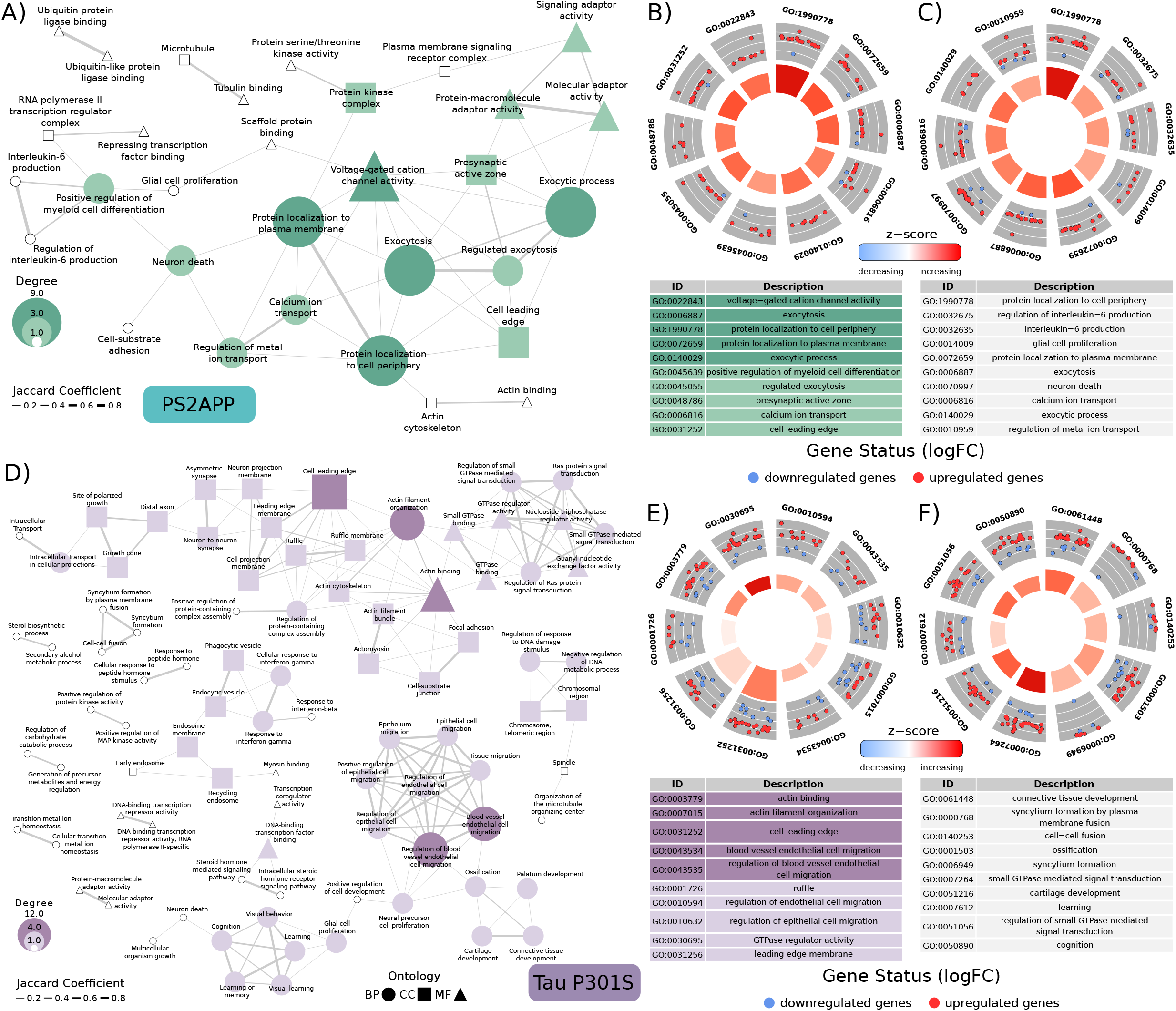
Gene Ontology network for biological processes (BP), cellular component (CC) and molecular function (MF). Gene Ontology (GO) term network of **(A)** PS2APP and **(D)** Tau P301S constructed using the proportion of overlapping genes between GO terms (Jaccard Coefficient > 0.15) as edges. Radial plots of 10 top GOBP terms ranked from the network by enrichment significance for **(B)** PS2APP and **(E)** Tau P301S. Radial plots of 10 top GO terms ranked from the network by connectivity (degree) for **(C)** PS2APP and **(F)** Tau P301S. Inner circle’s bar size maps significance of GO term enrichment and color maps the overall direction (upregulation or downregulation) of the term’s group of genes.

For the comparison of the Tau P301S mice *versus* WT controls, the functional enrichment analysis using DEGs revealed 95 terms (**Supplemental Table 2**), which include 56 BP, 25 CC and 14 MF enriched terms. The GO network for Tau P301S shows that the most connected terms are related to cytoskeleton dynamics such as “actin filament binding and organization” and “cell leading edge” (**Fig. 4D-E**). Among the most significantly enriched terms, we found terms such as “learning”, “cognition”, and “small GTPase mediated signal transduction” (**Fig. 4F**). Results of the functional enrichment analysis for both animal models are presented in **Supplemental Table 2**.

## 4. Discussion

Here, we evaluated the association of plasma GFAP levels with Aβ and tau pathologies in CU and CI individuals. We demonstrated that GFAP independently associates with Aβ, but not with tau pathology. To understand the impact of Aβ and tau pathology separately in GFAP-positive astrocyte phenotypes, we explored the molecular features of astrocytes isolated from the hippocampus of two AD mouse models (PS2APP and Tau P301S). Overall, we observed a limited overlap of DEGs between the Aβ and tau mouse models, indicating the unique nature of GFAP-positive astrocytic responses to different aspects of AD pathology. The findings presented in this study allowed the identification of functional categories and biological processes that are a distinctive response to Aβ or tau burden.

Astrocyte biomarkers have been increasingly investigated in AD, and they have already proved to be consistently altered in AD patients [6]. However, the production and consequent release of GFAP by astrocytes in response to AD pathological events seem heterogeneous. Specifically, while GFAP seems to be more associated with Aβ burden [8, 9, 24] it does not correlate with tau when corrected for Aβ pathology. On the other hand, studies investigating another astrocyte biomarker – the chitinase-3-like protein 1 (YKL-40) – demonstrated that YKL-40 seems to be more associated with tau pathology [25–27]. These findings highlight the importance of understanding how astrocytes react to specific aspects of AD pathology.

A large portion of the current knowledge regarding the astrocytic responses to AD pathology was obtained from *postmortem* studies in patients with dementia. At this point, the human brain is usually affected by several pathological components of AD simultaneously, hampering the understanding of the exact role each one is playing in the disease. The use of animal models provides a unique opportunity to explore isolated pathological features of AD and better comprehend its heterogeneous aspects. Our comparison of GFAP-positive astrocytes from animal models of Aβ and tau pathologies evidenced a scarce overlap of genes between these two transgenic lines. At the moment, only a few studies in the literature aimed at stratifying the unique astrocyte signatures triggered by Aβ or tau [5, 15]. Wu and colleagues specifically focused on the complement C1q cascade, demonstrating that tau pathology is enough to activate the classical complement pathway [15]. In a broader approach, Jiwaji et al. demonstrated a significant overlap of DEGs between astrocytes from Aβ and tau pathology. Our study and the one by Jiwaji and colleagues are highly complementary since present different methodologies. More specifically, studies used different brain regions (hippocampus *vs* whole cortex and spinal cord) and cell-sorting protocols that isolated distinct astrocyte populations (GFAP-positive astrocytes *vs* pan astrocytes) [5]. Despite the exciting insights provided by these recent studies, the understanding of astrocyte heterogeneity in AD is still in its early days, and further studies exploring region-specificity and disease/pathology stages are still needed.

Differential expression analysis in the PS2APP GFAP-positive astrocytes revealed that Aqp11 appeared as the most downregulated gene. In humans, AQP11 co-localizes to the endoplasmic reticulum (ER) and mitochondrial-associated membrane (MAM) to facilitate the transport of hydrogen peroxide [28, 29]. In the ER, AQP11-mediated peroxide transit seems to prevent toxicity during protein folding [29]. The downregulation of this peroxiporin in our analysis suggests the presence of ER stress in PS2APP GFAP-positive astrocytes. Additionally, as APP is synthesized in the ER, alterations in MAMs and ER stress are emerging as important pathological mechanisms of AD and interesting therapeutic targets to modify protein misfolding and synaptic failure [30–32]. Furthermore, the upregulation of ubiquitin-related genes (Uba52, Ubc) may indicate compensatory attempts to mitigate the results of ER stress. Ubiquitination is a major post-translational modification responsible for many actions in the cells, including protein degradation, kinases modification and regulation of signaling pathways. The PPI network analysis also evidenced that ER stress is an important alteration in PS2APP hippocampal astrocytes. In specific, the most connected proteins of this network (Ubc and Uba52) were associated with proteostasis. This functional category showed high linkage with others, especially intracellular trafficking, and DNA/RNA processing. Indeed, a recent systematic review of AD neuropathological literature showed proteostasis among the most consistent functional alterations of the disease, however its specificity for Aβ pathology was not demonstrated [33]. Furthermore, Tau P301S astrocytes PPI shows more genes associated with DNA/RNA processing function than PS2APP astrocytes. Site-specific phosphorylation regulates tau and, since nuclear tau protects the DNA [34, 35], its dysfunction can alter the chromatin landscape in pathological contexts [36, 37]. The DNA/RNA processing function contained Myc, Yy1, histone deacetylases, among other genes associated with DNA-binding and epigenetic GO terms. Although astrocyte and neuron chromatin dysfunction have been reported [38], it is not clear how tau-induced astrocyte dysfunction and associated chromatin alterations may influence other pathological features of AD.

The differential expression analysis of the Tau P301S astrocytes revealed dysregulation of insulin-related genes and downstream signaling kinases. Indeed, Ins1/2 and Pi3kca, proteins directly involved in regulating energetic brain metabolism, were highly connected and downregulated nodes in the Tau P301S PPI network. In this context, several pathophysiological links have been demonstrated between AD and metabolic disorders such as type 2 diabetes, obesity, and metabolic syndrome in humans and animal models [39–42]. Furthermore, it has been demonstrated that when insulin signaling is impaired in astrocytes, brain glucose uptake become less efficient, impairing CNS control of glucose homeostasis [42, 43]. As insulin plays a central role in regulating tau phosphorylation in the brain, deficiencies in insulin signaling could exacerbate neurodegeneration by promoting tau hyperphosphorylation in neurons [44, 45]. Despite *in vitro* studies demonstrating that insulin signaling in astrocytes is affected by Aβ exposure [46, 47], our findings showed that it does not seem to play a significant role in an animal model presenting Aβ deposition without tau pathology. In fact, the strong link between tau pathology and changes in insulin signaling has been reported in *postmortem* AD and other human tauopathies [48].

Interpreting transcriptomic changes at the pathway level is more consistent than single gene analysis, and it has the potential to offer insights into the biological processes disturbed in AD. Our functional enrichment analysis identified multiple biological processes driven by each specific aspect of AD pathology. Exocytosis-related processes are among the most significant biological processes enriched in the PS2APP mice. Interestingly, impaired exocytosis was already reported in AD, specifically in brain regions susceptible to neurodegeneration [49]. Corroborating, previous studies have been linking impaired exocytosis in the brain [50], and specifically in astrocytes [51], as a result of Aβ exposure, which might compromise proper neurotransmitter release. Astrocytes play a pivotal role in maintaining synaptic strength via vesicular exocytosis of glutamate and ATP [52]. Interestingly, our PPI analysis evidenced neurotransmission as one of the main functional categories disturbed in the PS2APP, but not in the Tau 301S astrocytes. In this sense, abnormalities in astrocyte exocytic activity might represent an underexplored aspect of AD pathology.

Neuroinflammation is now a well-recognized and active process detectable in the early stages of AD. A recent study using the [^11^C]PRB28 tracer revealed that inflammation and tau propagate jointly in the brain of AD patients in an Aβ-dependent manner [53]. We observed that abnormalities in inflammation-related processes were prominent in both Tau 301S and PS2APP astrocytes. Accordingly, it was previously demonstrated that Aβ and tau are able to independently trigger an inflammatory response that contributes to neurodegeneration in mice [54, 55], however, the presence of both pathologies simultaneously potentiates this process [54]. Additionally, the release of inflammatory mediators by astrocytes seems to be essential for Aβ-induced tau phosphorylation in primary neurons [56]. Although microglia are the main players in the brain immune response, astrocytes also respond to CNS threats by releasing a range of cytokines that perpetuate the harmful environment in AD [57–59].

## 5. Conclusion

The recent advances in understanding the neuropathological features of AD revealed a high degree of complexity in the astrocytic response to this disease. Additionally, the rapid growth in the AD fluid biomarkers field demonstrated that this complexity is reflected outside the brain. Indeed, comprehending AD blood biomarkers biological interpretation is one of the filed greatest challenges. The paucity of overlap between Aβ- and tau-driven DEGs strikingly shows the specific nature of GFAP-positive astrocyte responses to core AD pathology. Our results revealed multiple astrocyte-related biological processes and potential functional alterations triggered by each aspect of AD pathophysiology. Thus, developing novel biomarkers that reliably reflect AD heterogeneity seems critical to determine AD pathology’s underpinnings and consequently monitor the efficacy and target engagement of emerging anti-Aβ and anti-tau drugs and develop personalized treatment strategies.

## DECLARATIONS

### Ethics approval and consent to participate

Not applicable.

### Availability of data and materials

Datasets used in this study can be accessed via NCBI GEO portal (https://www.ncbi.nlm.nih.gov/geo/). Further intermediate data and codes generated are available from the corresponding author upon request.

### Author Contributions

MAB: conceptualization, data curation, formal analysis, data interpretation, figures, writing – original draft, and writing – review & editing.

BB: conceptualization, data curation, formal analysis, data interpretation, figures, writing – original draft, and writing – review & editing.

WSB: data curation, formal analysis, data interpretation, figures, writing – review & editing.

DGS: data curation, formal analysis, data interpretation, figures, writing – review & editing.

PCLF: data curation, formal analysis, data interpretation, figures, writing – review & editing. ASR: data curation, formal analysis, data interpretation, figures, writing – review & editing.

GP: data curation, formal analysis, data interpretation, figures, writing – review & editing.

JPF-S: data interpretation, writing – review & editing.

ALB: writing – review & editing.

NJA: writing – review & editing.

TKK: writing – review & editing.

HZ: writing – review & editing.

KB: writing – review & editing.

PR-N: writing – review & editing.

TAP: writing – review & editing.

ERZ: conceptualization, data interpretation, funding acquisition, supervision, writing – original draft, and writing – review & editing.

---

**Supplemental Figure 1.**
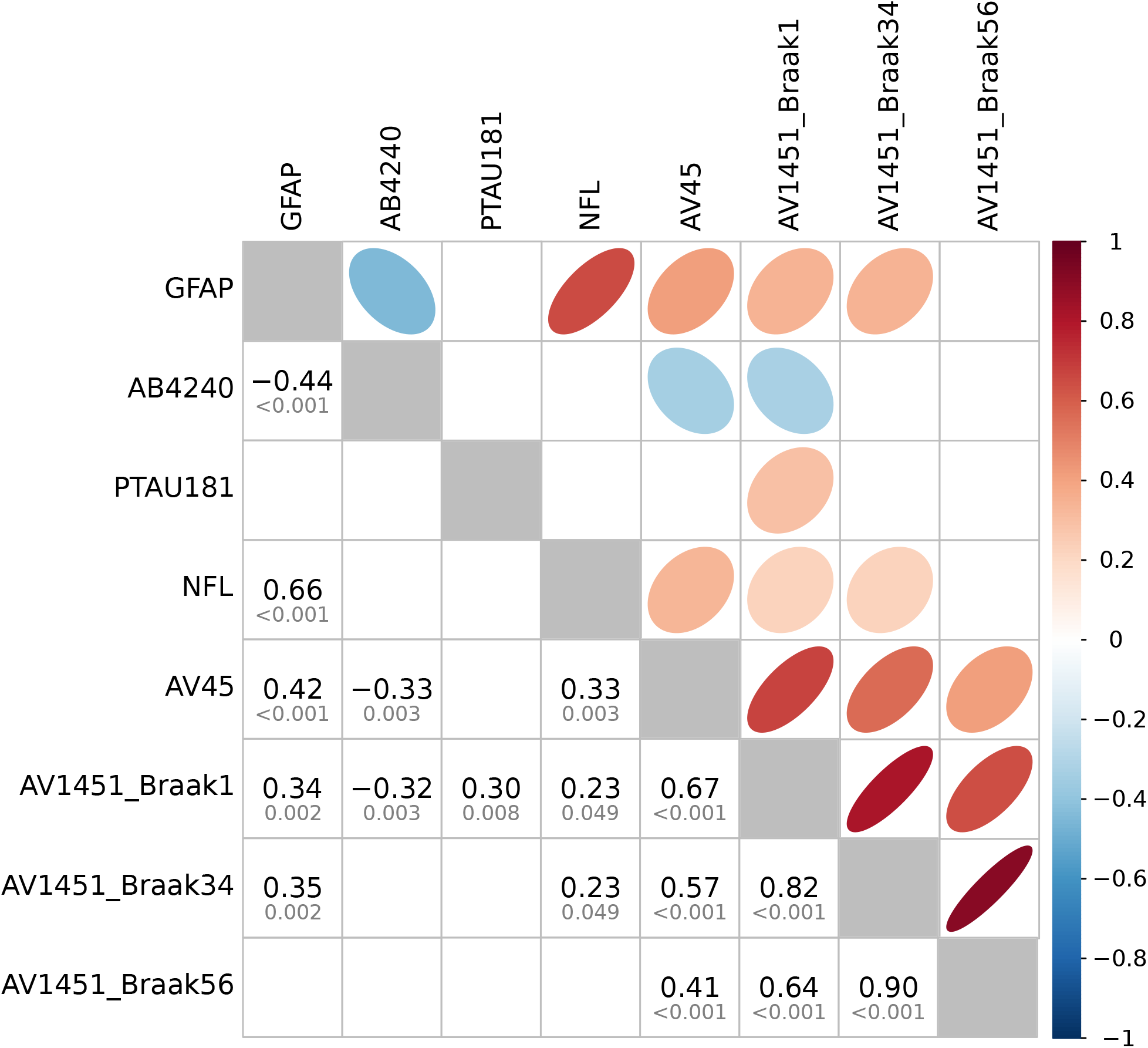
Correlation between GFAP and other AD biomarkers. Pairwise correlation matrix between GFAP and seven other known AD biomarkers. Pearson correlation was applied and FDR-adjusted (α = 0.05). The strength and direction of the correlation is represented by the color gradient in the upper triangle of the matrix. Coefficients (black) and adjusted p-values (gray) of the pairwise correlations are shown in the lower triangle.

**Supplemental Figure 2.**
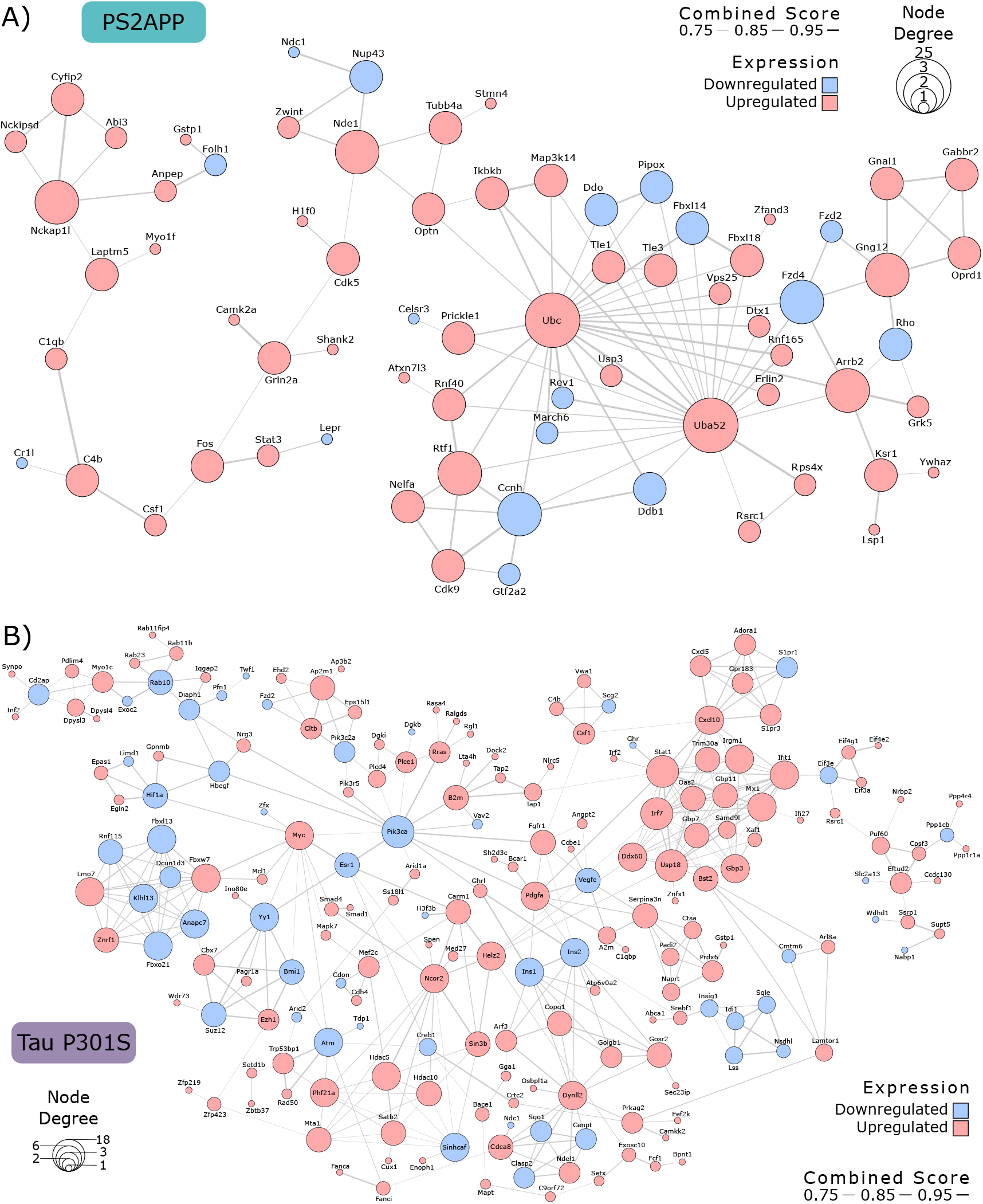
Regulation state of PS2APP and Tau P301S PPI astrocyte networks. Protein-protein interaction networks mapping regulatory states (up/down regulation) of differentially expressed genes in **(A)** PS2APP and **(B)** Tau P301S astrocytes. Combined score mapped to edges represents the combined confidence interaction scores provided by the STRING database for each pair of genes (values > 0.7 are considered high confidence scores).

